# Acetic acid enhances tolerance to long-term water deficit in tomato by partially buffering transcriptomic and proteomic reprogramming independently of canonical jasmonate signalling

**DOI:** 10.64898/2026.06.12.731858

**Authors:** Alberto Férez-Gómez, Lidia López-Serrano, Jesús Leal-López, Edurne Baroja-Férnandez, Goizeder Almagro, Antonio Gavira, Rafael Jorge León Morcillo, Javier Pozueta-Romero

## Abstract

Acetic acid (AA), a volatile compound present in diverse microbial-derived biostimulants, enhances drought tolerance in plants. In *Arabidopsis*, soil-applied AA action has been linked to histone H4 acetylation and activation of jasmonate (JA) signalling. However, the mechanisms underlying AA action in crops of agronomic interest remain poorly understood. Here, we used an integrative approach to evaluate the effects of soil-applied AA on fruit yield, physiological performance, and leaf transcriptomic and proteomic profiles of tomato plants grown under optimal and suboptimal irrigation conditions (OIC and SOIC, respectively). While AA had little effect under OIC, it significantly enhanced fruit yield and photosynthesis under SOIC. Long-term water deficit triggered extensive transcriptomic and proteomic reprogramming, particularly affecting photosynthesis, RNA processing, protein biosynthesis-, modification- and homeostasis-related processes. Under SOIC, AA induced marked molecular changes that were not consistent with activation of canonical JA signaling pathways. Notably, only ∼ 10% of the drought- or AA-responsive proteins were associated with corresponding transcript changes, highlighting a predominant role of regulatory layers beyond the transcriptional control to both long-term water deficit- and AA-induced protein remodeling. Strikingly, AA attenuated 47% and 35% of the transcriptomic and proteomic alterations induced by long-term water deficit, respectively. In addition, AA altered the abundance of numerous proteins that do not respond to drought, particularly ribosomal proteins and proteins involved in RNA processing. Collectively, our findings indicate that AA enhances tolerance to prolonged water deficit in tomato through mechanisms largely independent of canonical JA signaling and involving extensive downstream regulatory processes that partially mitigate stress-induced molecular reprogramming.

## INTRODUCTION

Water availability is the primary environmental stressor causing severe losses in global crop productivity (Ortiz-Bobea et al., 2021). Beyond the direct effects of drought, the environmental damage caused by intensive agricultural practices, including excessive fertilizer use and depletion of soil and water resources, threatens the sustainability of current production systems. Developing environmentally friendly strategies that improve crop productivity, and water and nutrient use efficiencies while reducing agrochemical inputs has therefore become a major challenge for modern agriculture (Penuelas et al., 2023; Maestre et al., 2025). Among these strategies, the use of biostimulants has emerged as a promising approach to enhance crop resilience under water-limited conditions.

Plants are metaorganisms that interact continuously with complex microbial communities through the exchange of chemical signals. Among these signals, microbial volatile compounds (VCs) act as semiochemicals mediating interkingdom communication (Schmidt et al., 2016; Schulz-Bohm et al., 2017). Air application of these compounds promotes growth and root developmental changes, enhances drought tolerance, and improves photosynthesis as well as nutrient and water acquisition (Gámez-Arcas et al., 2022). These physiological effects are associated with extensive transcriptomic, proteomic and redox-proteomic reprogramming involving proteostatic regulation of central metabolic pathways and long-distance root-to-shoot communication (Gámez-Arcas et al., 2022). We previously showed that soil application of fungal culture filtrates (CF) stimulates root growth and enhances fruit yield in pepper plants (Baroja-Fernández et al., 2021). These extracts contain VCs that, once distilled from the CFs and applied through irrigation, similarly increase fruit yield and root growth, suggesting that volatile constituents are major determinants of the response of plants to fungal CFs (Baroja-Fernández et al., 2021). To elucidate the biochemical and molecular mechanisms underlying CF action, we recently adopted a systems biology approach to characterize the response of tomato plants grown under optimal and suboptimal irrigation conditions (OIC and SOIC, respectively) to fungal CF application (López-Serrano et al., 2026). The data obtained provided evidence that CF improves long-term water deficit tolerance and fruit yield through novel regulatory mechanisms that involve mitigation of a substantial fraction of the stress-induced hormonal and transcriptional responses, thereby rendering plants less sensitive to water deficit (López-Serrano et al., 2026). However, whether these regulatory effects extend beyond the transcriptome to additional molecular layers, including translational and post-translational regulation, remains unknown. Moreover, the identity of the specific microbial VCs responsible for these effects has yet to be established.

Acetic acid (AA) is a microbial VC consistently detected in all biostimulant-active fungal CFs analyzed to date (Baroja-Fernández et al., 2021; Morcillo et al., 2022). This compound functions as a central signaling molecule and metabolic hub, with its involvement in protein acetylation representing a core mechanism by which plants regulate stress responses (Linster and Wirtz, 2018). Low-dose AA application through irrigation enhances drought tolerance in *Arabidopsis* and in several crops, including cassava, soybean, rice, cotton, apple and willow (Kim et al., 2017; Utsumi et al., 2019; Li et al., 2021; Ogawa et al., 2021; Rahman et al., 2021; Kong et al., 2022; Sun et al., 2022). AA treatment also enriches the populations of nitrogen-cycling, plant growth promoting microorganisms in the rhizosphere, thereby improving nitrogen uptake (Kong et al., 2022). Despite these observations, little is known about the molecular mechanisms underlying AA action. In Arabidopsis, AA induces histone H4 acetylation and primes jasmonate (JA) signaling as a mechanism to enhance drought tolerance (Kim et al., 2017). However, although several studies have described AA-induced transcriptomic responses under drought conditions (Kim et al., 2017; Utsumi et al., 2019; Sun et al., 2022), these responses have not been systematically compared with those elicited by water deficit itself. Furthermore, the proteomic effects of AA application remain completely unexplored. Consequently, it remains unclear whether AA confers drought tolerance by activating specific adaptive programs, by attenuating the molecular reprogramming induced by drought stress, or through a combination of both processes. To address these knowledge gaps, and to evaluate the contribution of AA to the enhanced tolerance to long-term water deficit previously observed following fungal CF application, here we have adopted an integrative approach encompassing fruit yield assessment, physiological characterization, and leaf transcriptomic and proteomic profiling in a commercial tomato hybrid grown in a semi-commercial greenhouse under OIC and SOIC, with or without soil application of AA. The results presented in this work uncover previously unrecognized regulatory mechanisms underlying AA action and demonstrate that AA alleviates the detrimental effects of long-term water stress partially by buffering stress-induced molecular reprogramming, providing valuable information for the development of sustainable agricultural practices.

## RESULTS

### Effect of soil-applied acetic acid on plant growth and fruit yield

Long-term water deficit markedly reduced vegetative growth and fruit productivity in tomato plants. Compared with OIC-grown plants, SOIC-grown plants displayed lower vegetative biomass production (**Figure 1A**), reduced number, yield and mean weight of marketable fruits, and a higher incidence of blossom-end rot (BER) (**Figure 1B**). Soil application of AA had little effect on leaf or stem biomass under either irrigation regime, but consistently promoted root growth (**Figure 1A**). Under SOIC, AA significantly improved reproductive performance by increasing the number and yield of marketable fruits by approximately 11%, while significantly reducing the number of non-marketable BER-affected fruits (**Figure 1B**). Mean fruit weight was not significantly altered by the treatment. Consequently, AA-treated plants required approximately 11% less water and nutrients to achieve the same marketable fruit yield under SOIC conditions (**Supplemental Figure 1**). Because BER is closely associated with calcium deficiency in fruits under irregular watering conditions (Topcu et al., 2022; Kabir and Díaz-Pérez, 2025), we quantified fruit calcium levels in all experimental groups. Long-term water deficit significantly reduced fruit calcium content, whereas AA treatment largely prevented this decline in SOIC-grown plants (**Figure 1C**).

**Figure 1:**
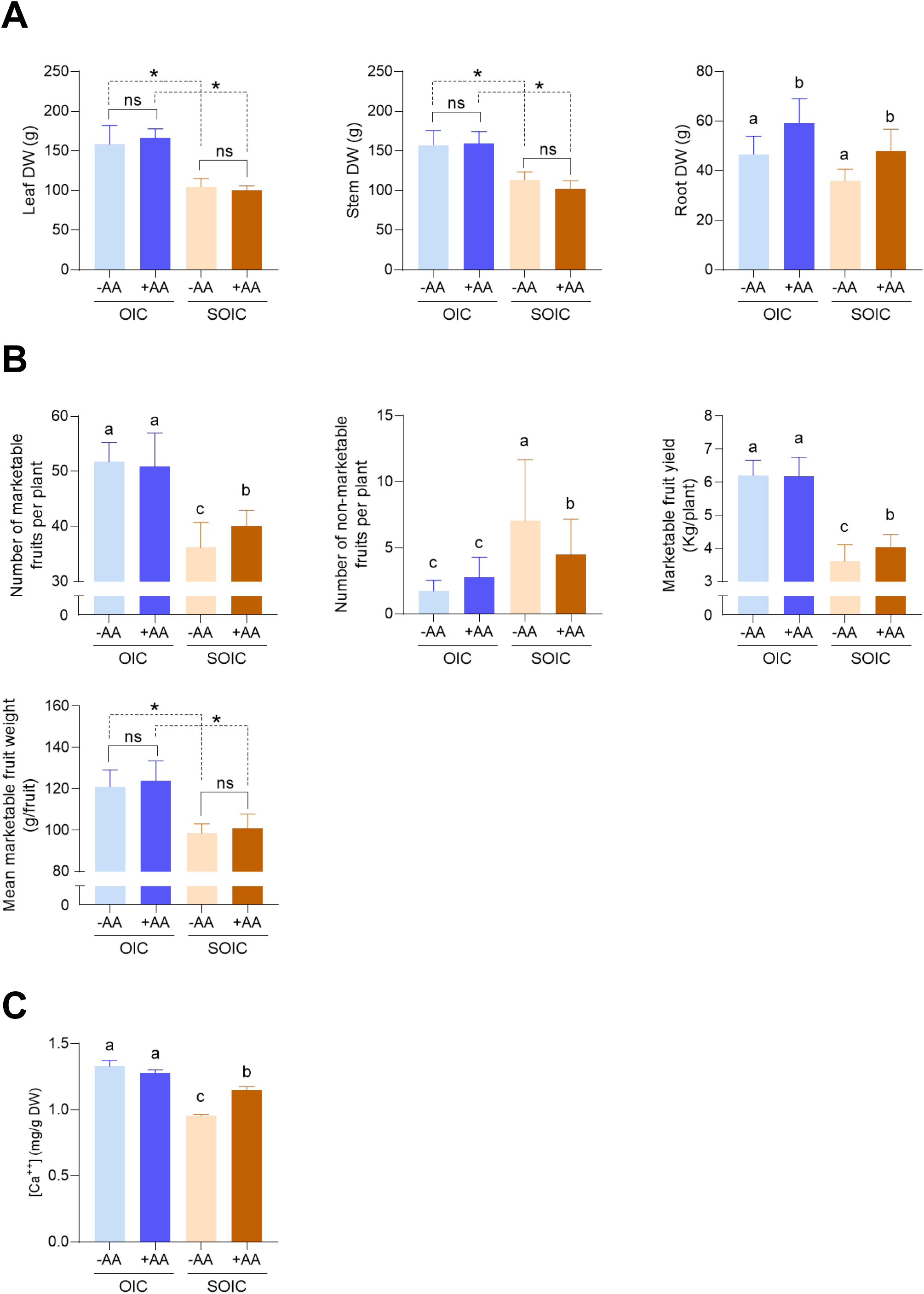
Effect of AA application on tomato plant growth and fruit yield. (A) Leaf, stem and root dry weights (DW) of plants grown under optimal and suboptimal irrigation conditions (OIC and SOIC, respectively), and with or without AA application. (B) Number of marketable and non-marketable fruits, and yield and mean weight of marketable fruits of plants grown under OIC or SOIC, with or without AA application. (C) Calcium content in marketable fruits of plants grown under OIC or SOIC, with or without AA application. Data were analysed by two-way ANOVA with irrigation (I) and AA treatment (T) as factors, followed by Fisheŕs LSD text (*P* < 0.05). When a significant I x T interaction was detected, different letters indicate significant differences among treatments. When no interaction was detected, but one factor was significant by a one-way ANOVA, significant differences were indicated with asterisks (*). Values represent the means ± standard deviations (SD) obtained from (A) 4 plants per irrigation condition and AA treatment and (B) 40 plants per irrigation condition and AA treatment, and (C) 3 fruits from 10 plants per irrigation condition and AA treatment.

### Effect of acetic acid application on photosynthesis, water potential and leaf composition

Long-term water deficit markedly impaired leaf gas exchange and plant water status. Compared with OIC-grown plants, plants grown under SOIC displayed significantly lower net photosynthetic rate (*A_n_*), stomatal conductance (*g_s_*), and transpiration rate (*E*) (**Figure 2A**). Notably, soil application of AA largely restored these parameters to levels comparable to those observed under OIC conditions. Consistent with these physiological improvements, midday leaf water potential (Ψ_md_) was significantly more negative in SOIC-grown plants than in OIC-grown plants, whereas AA treatment partially alleviated this decline under SOIC conditions (**Figure 2B**). Intrinsic water use efficiency (WUE*i*) increased under long-term water deficit, as expected from the stronger reduction in stomatal conductance relative to carbon assimilation. However, the AA treatment reduced the differences in WUE*i* between OIC- and SOIC-grown plants (**Figure 2B**), further indicating partial recovery of the physiological state of drought-stressed plants. Leaf amino acid and carbohydrate profiles differed markedly between irrigation regimes. In general, the AA treatment reduced the levels of sugars, including sucrose, glucose and fructose, as well as several of the most abundant amino acids, including Glu, Ser, Ala, GABA and Pro, in both OIC-and SOIC-grown plants (**Supplemental Table 1**).

**Figure 2:**
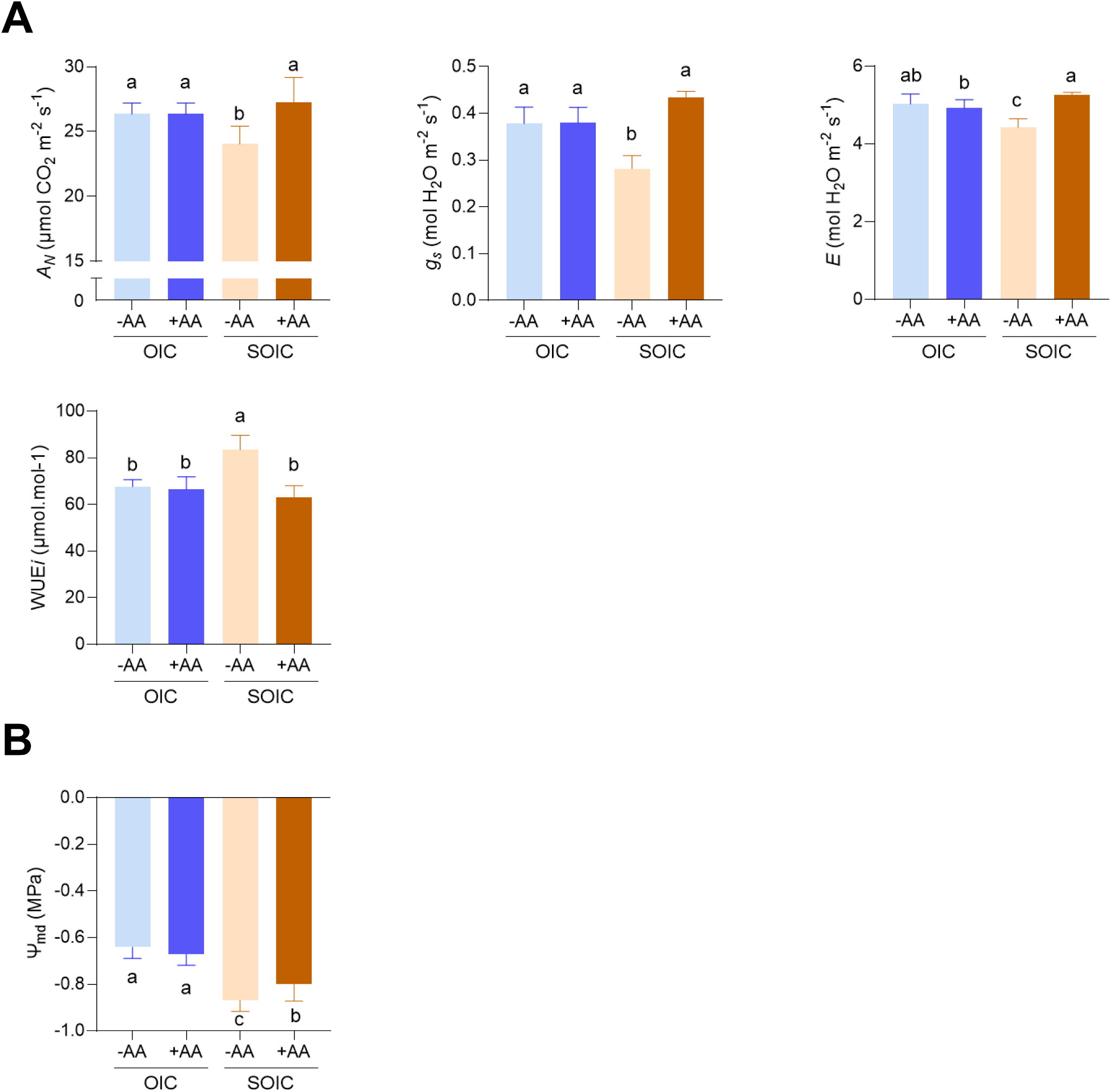
Effect of AA application on photosynthesis and water potential. (A) CO_2_ assimilation rate (*A_n_*), stomatal conductance (*g_s_*), transpiration (*E*) and intrinsic water use efficiency (WUE*i*) and (B) midday leaf water potentials (Ψ_md_) of plants grown under optimal and suboptimal irrigation conditions (OIC and SOIC, respectively), and with or without AA application. Different letters indicate significant differences at *P* < 0.05 (LSD test). Values represent the means ± standard deviations (SD) obtained from (A) 5 plants and (B) 10 plants.

### Effect of water stress and acetic acid on the leaf transcriptome and proteome

#### Effect of water stress and acetic acid application on the leaf transcriptome

RNA-seq analysis identified 24341 transcripts across all samples (**Supplemental Table 2**). Long-term water deficit induced extensive transcriptomic reprogramming, with 1131 transcripts upregulated and 1538 transcripts downregulated in leaves of SOIC-grown plants relative to OIC-grown plants (**Supplemental Table 3**). These 2669 transcripts were collectively defined as “drought-responsive transcripts” (**Supplemental Figure 2A**). Under SOIC conditions, the AA treatment also triggered major transcriptomic changes, increasing the abundance of 1393 transcripts decreasing that of 1019 transcripts (**Supplemental Table 4**). We refer to these 2412 transcripts as “AA-responsive transcripts” (**Supplemental Figure 2B**). Notably, a substantial fraction of the drought-promoted transcriptomic changes could be attenuated by AA application. Specifically, 384 of the 1131 transcripts that were upregulated by water scarcity, and 879 of the 1538 transcripts whose expression decreased due to this stress, exhibited opposite expression trends in SOIC-grown plants treated with AA (**Supplemental Table 5, Figure 3A**). Together, these 1263 transcripts represented 47% of all drought-responsive transcripts and were defined as “AA golden transcripts” because their drought-associated expression patterns were largely attenuated by AA treatment (**Figure 3B**).

**Figure 3:**
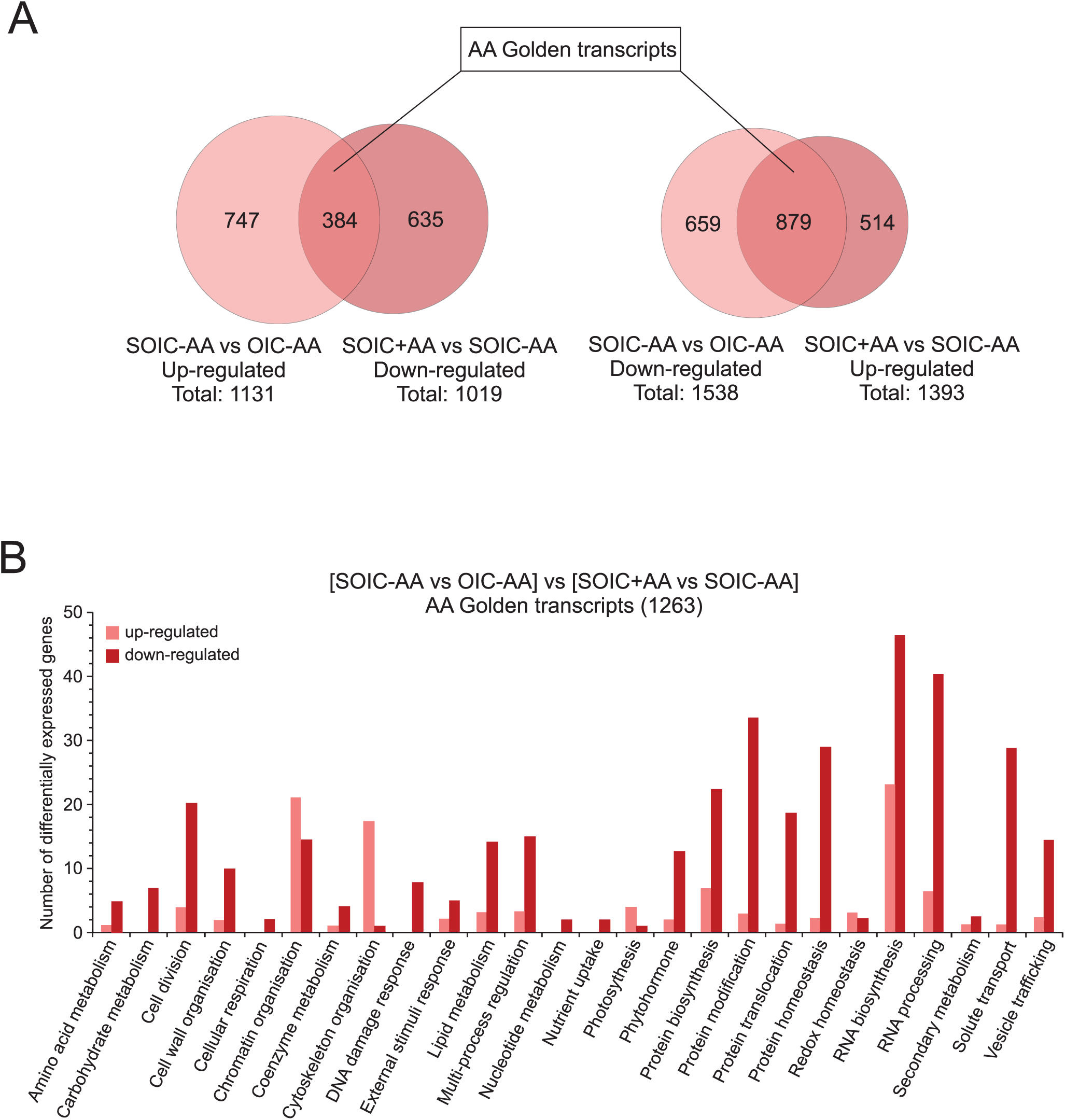
Identification of “AA golden transcripts”. (A) Venn diagram analysis illustrating the 1263 AA golden transcripts identified by comparing water deficit-driven transcriptomic changes (SOIC - AA vs. OIC - AA) with those promoted by AA in SOIC-grown plants (SOIC + AA vs. SOIC – AA). (B) Functional categorization of the “AA golden transcripts” according to Mapman (see also Supplemental Table 5). Numbers of up- and down-regulated transcripts in each categorical group are indicated by pale and deep red bars, respectively.

#### Effect of water stress and acetic acid application on the leaf proteome

The proteomic analysis identified 5931 proteins across all samples (**Supplemental Table 6**). Compared with OIC-grown plants, 105 proteins were up-regulated and 64 proteins were down-regulated in leaves of SOIC-grown plants (**Supplemental Table 7**). Functional classification of these 169 “drought-responsive proteins” using the Mapman tool revealed that long-term water deficit stress primarily reduced the levels of photosynthesis-related proteins, and increased those of secondary metabolism enzymes and stress-responsive processes operating beyond transcriptional control, including RNA processing, posttranslational protein modification, and protein homeostasis (**Supplemental Table 7, Figure 4A)**. AA treatment under SOIC conditions increased the levels of 44 proteins and decreased those of 99 proteins (**Supplemental Table 8, Figure 4B**). Functional analyses of these 143 “AA-responsive proteins” revealed that the AA treatment mainly caused a reduction in the levels of enzymes involved in secondary metabolism and stress-responsive proteins involved RNA processing, protein biosynthesis and posttranslational protein modification and homeostasis (**Supplemental Table 8**). Strikingly, AA counteracted a substantial proportion of the drought-induced proteomic changes. Specifically, 42 of the 105 proteins that were upregulated by long-term water scarcity, and 15 of the 64 proteins that were down-regulated by this stress, exhibited opposite accumulation trends in SOIC-grown plants treated with AA (**Supplemental Table 9, Figure 5A,B**). These 57 proteins, corresponding to 35% of all drought-responsive proteins, were defined as “AA-buffered drought-responsive proteins” or “AA golden proteins”. AA treatment under SOIC conditions also altered the abundance of 86 proteins whose levels were not significantly affected by water deficit (**Supplemental Table 10; Figure 5C**).

**Figure 4:**
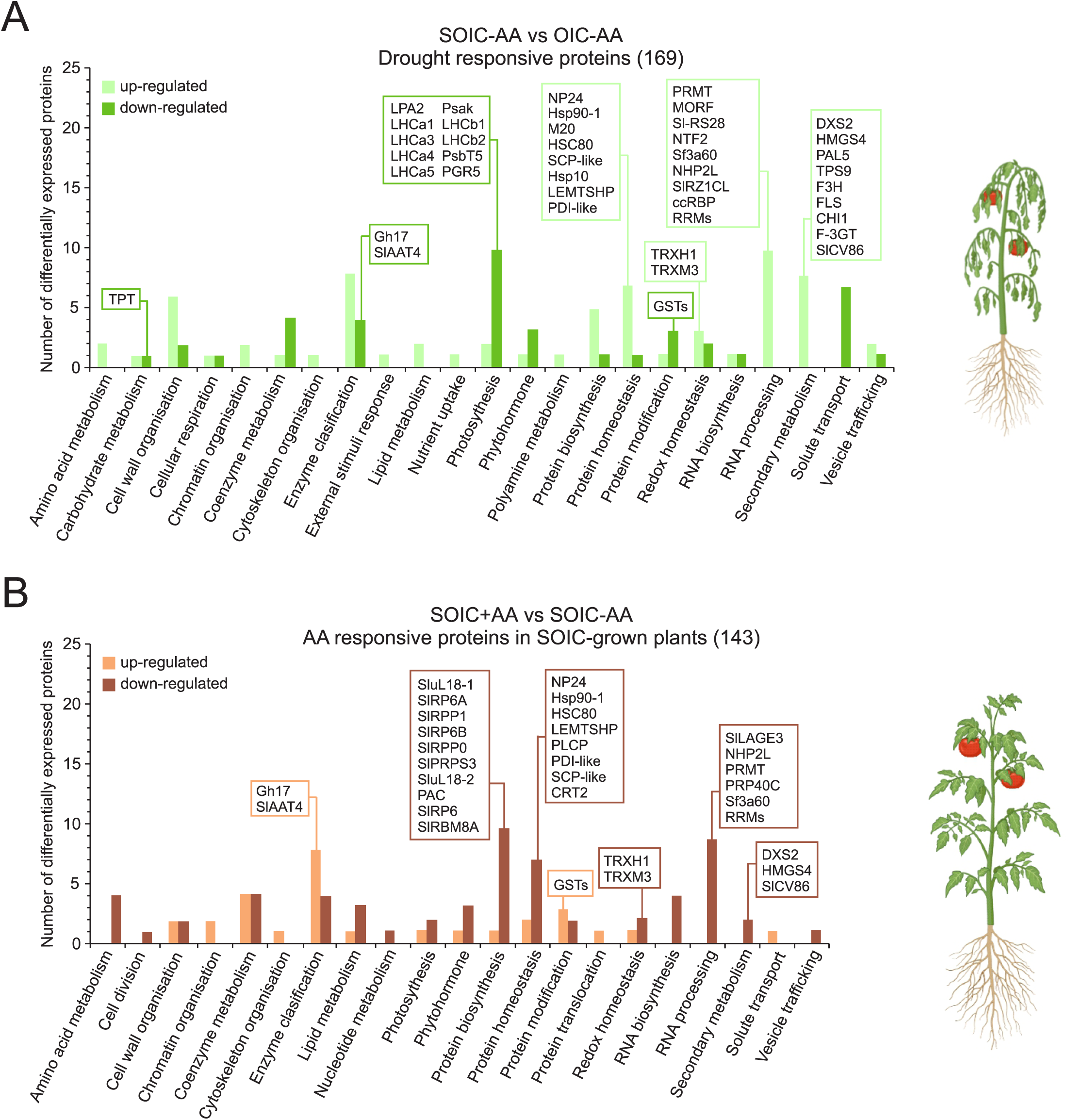
Proteome profiling of leaves of tomato plants grown under optimal and suboptimal irrigation conditions, with or without AA treatment. (A) Functional categorization of the 169 proteins differentially accumulated in leaves of tomato plants grown under suboptimal and optimal irrigation conditions without AA treatment (SOIC - AA and OIC - AA, respectively) (see also Supplemental Table 7). Numbers of up- and down-regulated proteins in each categorical group are indicated by pale and deep green bars, respectively. (B) Functional categorization of the 143 proteins differentially accumulated in leaves of plants grown under SOIC with or without AA treatment (SOIC + AA vs. SOIC - AA, respectively) (see also Supplemental Table 8). Numbers of up- and down-regulated proteins in each categorical group are indicated by pale and deep brown bars, respectively. Differentially accumulated proteins were sorted according to putative functional categories assigned by MapMan software and those discussed in the main text are boxed. The figure was created using BioRender.com.

**Figure 5:**
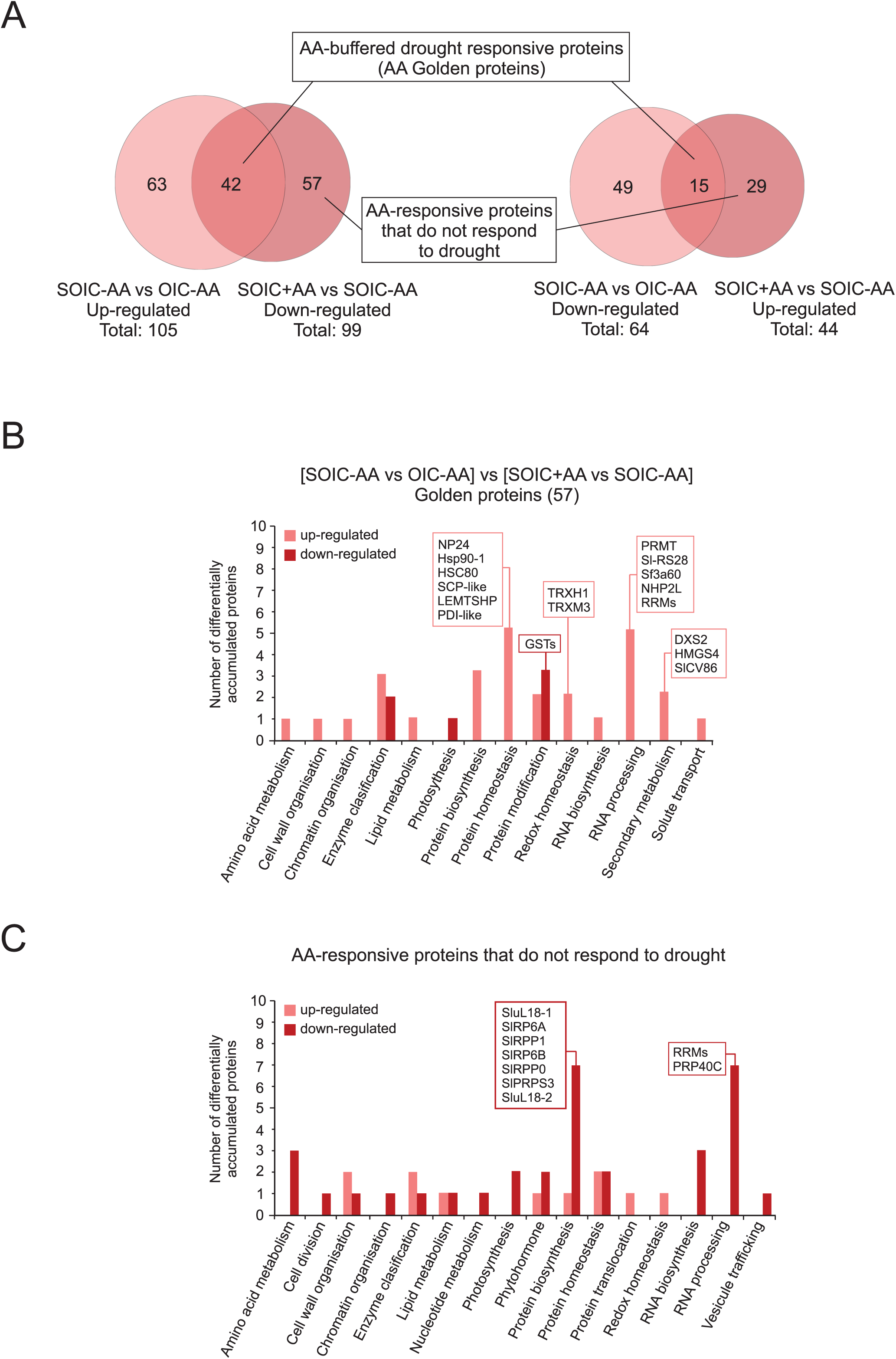
Identification of “AA golden proteins”. (A) Venn diagram analysis illustrating the 57 AA golden proteins identified by comparing water scarcity-driven proteomic changes (SOIC - AA vs. OIC - AA) with those promoted by AA in SOIC-grown plants (SOIC + AA vs. SOIC – AA). (B) Functional categorization of the “AA golden proteins” according to Mapman (see also Supplemental Table 9). Numbers of up- and down-regulated proteins in each categorical group are indicated by pale and deep red bars, respectively. Differentially expressed proteins discussed here are boxed.

## DISCUSSION

### AA is a major bioactive constituent of fungal CFs

Our results demonstrate that AA is sufficient to reproduce most of the beneficial effects previously exerted by AA-containing fungal CFs in tomato plants subjected to long-term water deficit (López-Serrano et al., 2026). Under SOIC, AA application improved photosynthetic performance and yield of marketable fruits, reduced the incidence of BER, and partially restored fruit calcium levels, whereas its effects under OIC were minimal. These findings strongly support the idea that AA is a major bioactive constituent of fungal CFs and further indicate that its biostimulant activity is strongly dependent on the physiological water status of the plant. AA-treated SOIC-grown plants also displayed less negative midday leaf water potential and enhanced transpiration rates relative to untreated SOIC-grown plants. Because calcium transport to developing fruits largely depends on the transpiration stream (Topcu et al., 2022; Kabir and Díaz-Pérez, 2025), the reduced BER incidence observed following AA treatment is likely associated with improved water flux and calcium allocation to fruits. The concomitant recovery of photosynthetic performance further supports the notion that AA alleviates the physiological constraints imposed by long-term water deficit. Importantly, AA application also reduced the water and nutrient input required to achieve equivalent yield of marketable fruits under SOIC conditions, highlighting its potential as an environmentally friendly biostimulant for sustainable tomato production under water-limited conditions. More broadly, the observation that AA exerts negligible effects under OIC but substantial benefits under SOIC suggests that its primary mode of action is not constitutive growth promotion, but rather attenuation of stress-associated physiological dysfunction.

### Proteomic responses to long-term water deficit and AA application are predominantly subjected to regulatory layers beyond the transcriptional control

A striking finding of this study is the limited correspondence between transcriptomic and proteomic responses to both long-term water deficit and AA application. Although plant responses to drought are generally considered to involve multi-layered regulatory mechanisms operating at transcriptional, post-transcriptional, translational and post-translational levels, previous analyses using leaves of tomato plants cv. M82 grown under highly controlled environmental conditions showed that 60% of the drought-responsive proteins were encoded by drought-responsive transcripts after 7 days of water deprivation (Liu et al., 2023). This indicated substantial transcriptional contribution to drought-induced proteome remodelling. In contrast, our data reveal a markedly different regulatory landscape in Macizo tomato plants grown under long-term water deficit in a semi-commercial greenhouse setting. Only 14% of the 169 drought-responsive proteins identified in this study were encoded by corresponding drought-responsive transcripts (**Supplemental Table 7**), indicating that the proteomic response to long-term water deficit is regulated predominantly downstream of transcript abundance. Consistent with this interpretation, a substantial proportion of the drought-responsive proteins identified in this study were functionally associated with RNA processing, protein synthesis, protein homeostasis, and post-translational modification (**Supplemental Table 7, Figure 4A**). Notably, none of the drought-responsive proteins identified in the present work overlapped with those (294) previously reported in drought-stressed M82 plants (Liu et al., 2023). This lack of overlap likely reflects the strong influence of genotype, developmental stage, stress duration and intensity, and environmental growth conditions on drought-response mechanisms, as shown in previous studies (Conti et al., 2019; Diouf et al., 2020).

A similarly limited transcript-protein correspondence was observed in the response to AA in SOIC-grown plants. Only 10% of the 143 AA-responsive proteins identified in this study were encoded by AA-responsive transcripts (**Supplemental Table 8**), indicating that AA-induced proteome reprogramming also occurs largely through regulatory layers beyond transcriptional control. In agreement with this view, approximately one third of the AA-responsive proteins are involved in RNA processing and protein synthesis and protein homeostasis-related processes (**Supplemental Table 8, Figure 4B**). Collectively, these findings indicate that both chronic water deficit and AA application primarily reshape the leaf proteome through regulatory layers downstream of transcription. More broadly, they suggest that the physiological and molecular effects of AA are primarily mediated not by large-scale transcriptional activation, but by modulation of downstream regulatory processes that influence proteome stability and cellular homeostasis under stress conditions, emphasizing the need to integrate proteomic analyses when dissecting plant stress responses.

### Long-term water deficit promotes extensive transcriptomic and proteomic reprogramming that do not involve activation of the AA biosynthetic and canonical JA signaling pathways

Previous studies in Arabidopsis grown under tightly controlled environmental conditions showed that drought stress reduces the expression of glycolytic genes while increasing that of enzymes involved in AA biosynthesis, thereby promoting histone H4 acetylation and activation of JA signalling as a mechanism to enhance drought tolerance (Kim et al., 2017). In contrast, our transcriptomic and proteomic analyses carried out in this work using tomato plants grown under semi-commercial greenhouse conditions did not reveal significant drought-induced changes in the expression of glycolytic and AA biosynthetic pathways (**Supplemental Table 3, Supplemental Table 7**). Moreover, inspection of the 2669 drought-responsive transcripts identified in this study failed to uncover a prominent signature of canonical JA signalling. Although a limited number of genes associated with oxylipin metabolism and wound responses were detected, core components of the JA biosynthetic and signalling machinery were largely absent. Instead, drought-responsive transcripts included numerous genes associated with ABA, ethylene, auxin and, to a lesser extent, cytokinin signalling. These observations indicate that the molecular mechanisms underlying acclimation to long-term water deficit in tomato differ substantially from those described in Arabidopsis and are strongly influenced by species, developmental stage, stress duration and intensity, and growth conditions.

Long-term water scarcity induced pronounced changes in the levels of proteins with well-established roles in drought responses. Leaves of SOIC-grown plants accumulated markedly lower levels of proteins involved in photosynthesis than those of OIC-grown plants (**Supplemental Table 7, Figure 4A**). These included core components of the photosystem complexes (particularly light harvesting proteins) and the plastidial triose phosphate/phosphate translocator (TPT), which mediates the export of Calvin-Benson cycle intermediates from the chloroplast to the cytosol. Moreover, leaves of SOIC-grown plants accumulated higher levels of TRXH and TRXM than those of OIC-grown plants (**Supplemental Table 7, Figure 4A**). Under different stress conditions, these thioredoxins typically accumulate to regulate the redox status-dependent activity of proteins involved in stomatal aperture and photosynthesis, and in maintaining the critical balance between oxidizing and reducing agents and the active structure of proteins (Park et al., 2009; Sahrawy et al., 2022; Zhai et al., 2022; Cui et al., 2025). Collectively, these alterations in protein levels provide a mechanistic explanation for the reduced stomatal aperture, decreased photosynthetic performance, and consequently reduced growth and fruit yield observed in SOIC-grown plants.

Long-term water deficit also promoted the accumulation of numerous proteins involved in RNA processing (particularly RNA recognition motif (RRM) domain proteins) and protein homeostasis (e.g. LEMTSHP, NP24, M20, Hsp90-1, HSC80, Hsp10) that play important protective roles in responses to environmental stress factors (Goel et al., 2010; Bashir et al., 2020; Wei et al., 2020; Sato et al., 2024) (**Supplemental Table 7, Figure 4A**). These responses further support the notion that acclimation to chronic water deficit relies heavily on regulatory processes operating downstream of transcription. Interestingly, drought stress also increased the abundance of 1-deoxy-D-xylulose-5-phosphate synthase (DXS) and hydroxymethylglutaryl-CoA synthase (HMGS) (**Supplemental Table 7, Figure 4A)**, which are key rate-limiting enzymes in the synthesis of isoprenoid compounds. Because isoprenoids are essential for photosynthesis, growth, and stress acclimation (Pulido et al., 2012), increased accumulation of these enzymes may reflect an attempt to sustain metabolic homeostasis under adverse conditions. However, DXS is subject to strict proteostatic regulation and tends to accumulate as inactive aggregates during stress (Rodríguez-Concepción et al., 2019). Thus, the increased abundance of DXS observed in SOIC-grown plants may not necessarily reflect enhanced enzymatic activity and isoprenoid biosynthesis, but rather accumulation of inactive or misfolded protein forms under chronic drought conditions.

### Exogenous AA application attenuates drought-induced transcriptomic and proteomic reprogramming independently of canonical JA signalling

Previous studies in Arabidopsis proposed that AA-mediated drought tolerance relies on histone H4 acetylation-dependent activation of JA biosynthesis and signalling pathways (Kim et al., 2023). Enhanced JA accumulation and signalling promoted by AA application have also been reported in drought-stressed cotton, apple, and willow plants (Li et al., 2021; Kong et al., 2022; Sun et al., 2022), whereas no such effects were observed in cassava (Utsumi et al., 2019). Inspection of the 2412 AA-responsive transcripts identified in this study revealed only a limited number of genes potentially associated with JA metabolism, whereas core components of the canonical JA biosynthetic and signalling machinery were largely absent (**Supplemental Table 4**). In contrast, genes related to auxin, cytokinin, ABA and ethylene signalling were more prominently represented among AA-responsive transcripts. These observations indicate that the mechanisms underlying AA-mediated drought tolerance in tomato differ substantially from the canonical JA-associated model proposed in Arabidopsis, and highlight the strong influence of species-specific and environmental factors on AA responses.

We previously showed that fungal CFs attenuate a large subset of drought-responsive transcriptional changes in tomato, defining a group of “CF golden transcripts” associated with improved drought tolerance (López-Serrano et al., 2026). Similarly, soil AA application mitigated or prevented the changes in the expression of 47% of the drought-responsive transcripts identified in this study (**Supplemental Table 5**). Notably, approximately 20% of the previously identified 638 “CF golden transcripts” overlapped with the “AA golden transcripts” identified here (**Supplemental Table 11**), suggesting the existence of a core set of drought stress-responsive genes particularly sensitive to AA-containing microbial-derived biostimulants. A similar buffering effect of AA was observed at the proteomic level. AA mitigated approximately 35% of the proteomic alterations induced by long-term water deficit, including changes affecting proteins involved in secondary metabolism, isoprenoid biosynthesis, redox regulation, RNA processing, and protein homeostasis and redox modification (**Supplemental Table 9, Figure 5B**). Among these “AA golden proteins” were DXS and HMGS, thioredoxins such as TRXH and TRXM, and several molecular chaperones including Hsp90-1 and HSC80. Because many of these proteins are associated with photosynthetic acclimation, proteostasis, and stress-related metabolic adjustments, attenuation of their drought-induced accumulation provides a mechanistic explanation for the improved stomatal conductance, photosynthetic performance and, consequently, improved fruit yield conferred by AA in SOIC-grown plants (**Figure 6**).

**Figure 6:**
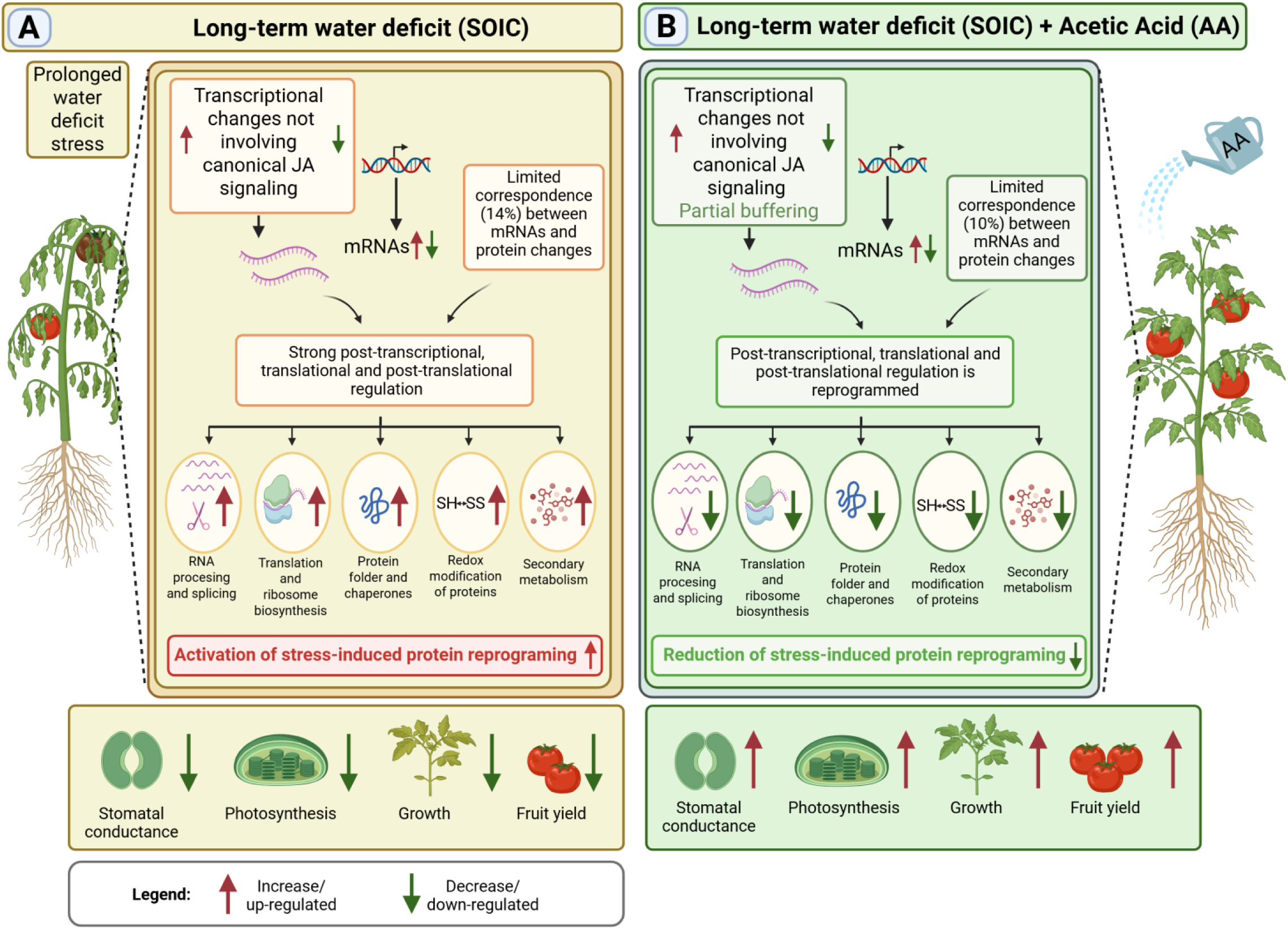
Proposed mode of action of acetic acid in SOIC-grown tomato plants. **Left panel:** Long-term water deficit reduces stomatal conductance, photosynthetic CO₂ fixation, vegetative growth, and fruit yield. These physiological alterations are associated with extensive transcriptomic and proteomic reprogramming. Despite the large number of drought-responsive transcripts and proteins, only a small proportion of drought-responsive proteins are encoded by drought-responsive transcripts. This observation, together with the accumulation of proteins involved in RNA processing, protein synthesis, protein folding, protein homeostasis, and redox regulation, suggests that acclimation to chronic water deficit is governed largely by post-transcriptional, translational, and post-translational regulatory mechanisms. The resulting activation of stress-induced protein reprogramming contributes to the establishment of the drought-stressed physiological state. **Right panel:** Application of AA through the irrigation water enhances stomatal conductance, photosynthetic CO₂ fixation, growth, fruit yield, and fruit quality under long-term water deficit conditions. As observed for drought responses, correspondence between AA-responsive transcripts and proteins is limited, indicating that AA-induced molecular reprogramming also operates predominantly through regulatory layers downstream of transcription. According to the proposed model, AA attenuates a substantial fraction of the transcriptomic and proteomic changes induced by long-term water deficit, thereby reducing the extent of stress-induced molecular reprogramming and maintaining plants in a physiological and molecular state closer to that of optimally irrigated plants. In addition, AA promotes specific proteomic adjustments that are independent of the drought response itself, including changes affecting proteins associated with RNA processing, translation-related functions, protein homeostasis, and redox regulation. Collectively, these effects contribute to improved photosynthetic performance, growth, fruit yield, and overall tolerance to prolonged water deficit. The figure was created using BioRender.com.

Interestingly, AA also affected the abundance of numerous proteins whose levels were not significantly altered by water deficit, including several functionally important ribosomal proteins and proteins involved in RNA processing (**Supplemental Table 10, Figure 5C**). Together with the remarkably low transcript–protein correspondence observed throughout this study, these findings suggest that AA does not simply attenuate drought-induced molecular responses, but also promotes selective remodelling of regulatory processes associated with RNA metabolism, translation-related functions and proteostasis (**Figure 6**). This interpretation is consistent with the predominance of AA-responsive proteins involved in RNA processing, protein synthesis and protein homeostasis, and reinforces the notion that the beneficial effects of AA operate largely through regulatory layers downstream of transcription.

### Additional and concluding remarks

Our findings support a model in which exogenous AA alleviates the detrimental effects of prolonged water deficit in tomato not through activation of large-scale defense programs, but through partial attenuation of drought-induced molecular and physiological reprogramming combined with additional regulatory adjustments affecting translation-related processes (**Figure 6**). AA treatment mitigated extensive drought-induced transcriptomic and proteomic reprogramming, particularly affecting proteins associated with photosynthesis, stomatal regulation, RNA metabolism, and protein homeostasis and redox modification. As a consequence, AA-treated plants remained physiologically and molecularly closer to plants grown under optimal irrigation conditions. At the same time, AA also promoted proteomic alterations that were independent of the drought response itself, including downregulation of multiple ribosomal proteins and proteins involved in RNA processing. These observations indicate that AA does not simply reverse the drought-induced molecular state, but actively reshapes cellular regulatory and metabolic processes under chronic water-limited conditions. This mode of action differs fundamentally from the transcriptionally driven mechanism previously proposed in Arabidopsis which involves histone H4 acetylation-dependent activation of JA signaling, further underscoring the strong context dependency of AA-mediated stress responses across plant species and environmental conditions.

The molecular mechanisms underlying the proteomic remodeling promoted by AA remain to be fully elucidated. We must emphasize that, beyond the well-established role of histone acetylation in chromatin structure and transcriptional regulation, acetylation of non-histone proteins has emerged as a major regulator of protein stability, enzymatic activity, protein–protein interactions, subcellular localization, and crosstalk with other post-translational modifications (Drazic et al., 2016; Wang et al., 2021). Protein acetylation is therefore increasingly recognized as an important component of plant acclimation to environmental stress, including drought (Linster et al., 2015; Linster and Wirtz, 2018; Guo et al., 2022). In this context, it is particularly noteworthy that several ribosomal proteins and heat-shock proteins differentially affected by AA treatment, including multiple 60S and 40S ribosomal subunits and Hsp90-related proteins (**Supplemental Table 8, Figure 4B**), belong to protein families that are frequently subject to non-histone acetylation (Oh et al., 2017; Xia et al., 2022). Because AA availability directly influences intracellular acetyl-CoA pools (Liu et al., 2018; Fu et al., 2020), it is conceivable that AA-mediated stress attenuation partly involves altered acetylation dynamics of proteins associated with translation, proteostasis, and RNA metabolism. Particularly intriguing is the observation that AA reduced the abundance of several functionally important ribosomal proteins that do not respond to drought, including two paralogs of the regulatory protein S6, the acidic stalk proteins P0 and P1, and two uL18 large-subunit proteins. Together with the concomitant reduction of proteins involved in RNA processing, these changes suggest that AA promotes selective remodeling of ribosome-associated functions and RNA metabolism rather than a global suppression of protein synthesis. Although this hypothesis remains speculative, it provides a plausible mechanistic framework linking AA metabolism to the extensive downstream regulatory effects observed in this study.

The precise contribution of individual AA-buffered drought-responsive proteins (AA golden proteins) and drought-independent AA-responsive proteins to enhanced drought tolerance will require further investigation using transgenic and gene-editing approaches. Future studies should also determine whether AA-mediated buffering extends to additional regulatory layers beyond the transcriptome and proteome. Importantly, exogenous AA application has been reported to enhance tolerance to multiple abiotic and biotic stresses (Chen et al., 2019; Rahman et al., 2019), raising the possibility that AA functions as a broad-spectrum modulator of stress-state transitions in plants. More broadly, our results suggest that comparative identification of stress-buffered “golden proteins” by contrasting stress-induced molecular responses with those elicited by AA application under stress conditions represents a powerful strategy to identify regulatory nodes associated with stress resilience. Such approaches may facilitate the discovery of molecular biomarkers of stress buffering and support the development of environmentally sustainable strategies aimed at improving crop performance under adverse environmental conditions.

## MATERIAL AND METHODS

### Experimental site, plant material, growth conditions and sampling

Assays were conducted between March and July 2023 in a semi-commercial polyethylene greenhouse located at the “Instituto de Hortofruticultura Subtropical y Mediterránea” in Algarrobo (Málaga, Spain; 36°45’32.0“N 4°02’28.7“W). Maximum midday greenhouse air temperature ranged between 32.6 and 41.2 °C, with a minimum night temperature ranging between 6.2 and 16.2 °C (**Supplemental Figure 3A**). Maximum relative humidity ranged between 84.8 and 94.9 %, whereas minimum relative humidity ranged between 13.7 and 28.9 % (**Supplemental Figure 3B**). Values of photosynthetic active radiation (PAR, 400-700 nm) recorded at midday were 534 ± 111 µmol m^−2^ s^−1^ (**Supplemental Figure 3**).

Plants of the commercial tomato hybrid Macizo F1 (Ramiro Arnedo Semillas) were grown essentially as described in López-Serrano et al. (2026). At the 2–4 true-leaf stage, plants were assigned to one of two irrigation regimes: OIC or SOIC (**Supplemental Figure 4**), the latter receiving 60% of the irrigation volume supplied to OIC-grown plants. Irrigation frequency and nutrient solution volume were adjusted according to substrate volumetric water content monitored using TEROS 12 sensors (METER Group, USA) (**Supplemental Figure 5A**). Substrate temperature (**Supplemental Figure 5B**) and bulk electrical conductivity (BEC) (**Supplemental Figure 5C)** were also recorded every 15 min using TEROS 12 sensors. AA treatments were applied through the irrigation system once per week from the 2–4 true-leaf stage until the onset of fruit harvest, resulting in a total of 14 applications (**Supplemental Figure 4**). For each application, pure AA was added to the nutrient solution to a final concentration of 20 mM. The AA-containing nutrient solution was supplied during a 5-h irrigation period between 08:00 and 13:00 h. Sampling of leaves at the 42 days after water shortage (DAWS) time point, and fruit harvesting were conducted as described in López-Serrano et al. (2026) using 40 plants per irrigation and AA treatment randomly set throughout the greenhouse. Fruits were classified into “marketable” and “non-marketable” according to López-Serrano et al. (2026). The agronomic water use efficiency for yield (WUEyld) was calculated as the ratio between the amount of irrigation water supplied and the total marketable fruit production per plant. Biomass studies were conducted as described in López-Serrano et al. (2026) using 4 plants per irrigation condition and AA treatment harvested at the end of the campaign (**Supplemental Figure 4**).

### Determination of gas exchange parameters

Measurements of net rates of CO_2_ assimilation (*A_n_*), stomatal conductance (*g_s_*) and transpiration (*E*) values were conducted between 12.00 and 13.30 pm (GMT) using the portable photosynthesis system LCpro-SD (ADC BioScientific Ltd.) and young leaves (the 8^th^-9^th^ leaf from the apex) of 5 plants per treatment and irrigation condition at the 65 DAWS time point (**Supplemental Figure 4**). Values of environmental parameters inside the leaf chamber were 1000 µmol m^−2^ s^−1^ PAR, 25 °C air temperature and 400 ppm air CO_2_. Values of intrinsic water use efficiency (WUE*i*) were calculated as the ratios of *A_n_* to *g_s_*, and *A_n_* to *E*, respectively, as described by Flexas et al. (2016).

### Biochemical characterization

Levels of soluble sugars and amino acids in leaves at the 42 DAWS time point were determined as described in Baroja-Fernández et al. (2021). Fruit calcium content was quantified at the ionomic service of Centro de Edafología y Biología Aplicada del Segura (CEBAS, Murcia, Spain) by ICP emission spectrometry (iCAP 6000, Thermo Scientific, Cambridge, England). For each irrigation condition and AA treatment, values were obtained from 3 pooled samples, each pool consisting of leaf material collected from 10 individual plants.

### Transcriptomic analysis, validation of data and identification of hormone-related genes

Transcriptomic analyses of leaves at the 42 DAWS time point were conducted essentially as described in López-Serrano et al. (2026), except that RNA sequencing was carried out at the Centre Nacional d’anàlisi genómica (CNAG). For each irrigation condition and CF treatment, RNA was extracted from three pooled samples, each pool consisting of material collected from 10 individual plants. Validation of the results of the RNA-seq data was conducted by RT-qPCR as indicated in Sánchez-López et al. (2026) using primers listed in **Supplemental Table 12**. To assess the potential involvement of plant hormone signalling pathways in the responses to long- term water deficit and AA treatment, drought- and AA-responsive transcripts were manually inspected based on their functional annotations. Gene annotations were obtained from the Sol Genomics Network (SGN; Fernandez-Pozo et al., 2015), UniProt Knowledgebase (UniProt Consortium, 2025), Plant Reactome (Naithani et al., 2020), and previously published literature on hormone signalling pathways in tomato and Arabidopsis. Genes were classified as potentially associated with JA, ABA, auxin, cytokinin or ethylene signalling when their annotations indicated established roles in hormone biosynthesis, perception, signal transduction, transcriptional regulation, or downstream hormone-responsive processes. Particular attention was paid to the identification of genes belonging to the canonical JA pathway, including genes involved in JA biosynthesis (e.g. lipoxygenases, allene oxide synthases, allene oxide cyclases and 12-oxophytodienoate reductases), JA perception and signalling (e.g. COI1, JAZ and MYC family proteins), and JA-responsive defence-related genes such as proteinase inhibitors. Similar criteria were applied to identify genes associated with ABA, auxin, cytokinin and ethylene signalling pathways.

### Proteomic analysis

High-throughput, isobaric labelling-based differential proteomic analyses of leaves were conducted at the Proteomic Facility at the Centro Nacional de Biotecnología (CNB-CSIC) essentially as described in Gámez-Arcas et al. (2022b), but the tryptic peptides were labelled using TMT-12plex Isobaric Mass Tagging kit (Thermo Fischer Scientific). The cut-off for identifying differentially accumulated proteins was established at a FDR ≤ 0.05% and log2 ratios (+ treatment vs. - treatment) > 0.3 (for proteins whose expression was up-regulated by the treatment) or < −0.3 (for proteins whose expression was down-regulated by the treatment).

### Statistical analyses

Statistical analyses were performed using R Core Team statistical software (https://www.R-project.org) within the RStudio integrated development environment (http://www.posit.co). Data were analysed using a two-way Analysis of Variance (ANOVA), implemented with the aov function from the stats package in R (v4.3.1; R Core Team 2023), where AA application treatments (Treatment, T) and irrigation conditions (Irrigation, I) were considered as fixed factors of the analysis. Statistical significance was accepted at *P*<0.05. When a significant interaction between Treatment and Irrigation was detected, a one-way ANOVA was performed using the combined factor (T x I). When double interaction was not found in the two-way ANOVA, but one or both individual factors were statistically significant, a one-way ANOVA was performed taking into consideration only the significant factor. Specifically, for AA Treatment effects, -AA and +AA conditions were compared within the same irrigation regime, whereas for the Irrigation effects, OIC and SOIC were compared within the same treatment type. Means were compared using Fisher’s least significant difference (LSD) test at *P<*0.05 with the LSD.test function from the agricolae package (https://CRAN.R-project.org/package=agricolae).

## ACKNOWLEDGEMENTS

This research was funded by the Ministerio de Ciencia, Innovación y Universidades (MCIU), Agencia Estatal de Investigación (AEI) / 10.13039/501100011033/ (grant PID2022-137292NB-I00) and European Union Next Generation EU/PRTR (grant TED2021-130603B-C21). L L-S is beneficiary of a Juan de la Cierva 2022 postdoctoral fellowship (reference number JDC2022-049385-I) funded by MCIN/AEI /10.13039/501100011033 and European Union Next Generation EU/PRTR. J L-L acknowledges a pre-doctoral fellowship funded by MCIU/AEI/10.13039/501100011033 and by “ESF Investing in your future”. We thank José Manuel Ramos Martín, Sara Sánchez Segovia, Gonzalo González Gil, Juan Francisco Ruiz Solanilla and Laura Frías for technical support and conducting experiments at the IHSM greenhouse. We also thank Anna Esteve-Codina and Marta Gut (both from the Centre Nacional d’anàlisi genómica, CNAG) for their help with the the RNA-seq studies.

## SUPPLEMENTAL FIGURES

**Supplemental Figure 1: Agronomic water use efficiency (WUEyld) of tomato plants grown under optimal and suboptimal irrigation conditions (OIC and SOIC, respectively) and with or without AA treatment.** Values represent the means ± SD obtained from 40 plants.

**Supplemental Figure 2: Transcriptome profiling of leaves of tomato plants grown under OIC and SOIC, with or without AA treatment.** (A) Functional categorization of the transcripts differentially expressed in leaves of tomato plants grown under suboptimal and optimal irrigation conditions without AA treatment (SOIC-AA and OIC-AA, respectively) (see also **Supplemental Table 3**). Numbers of up- and down-regulated transcripts in each categorical group are indicated by pale and deep green bars, respectively. (B) Functional categorization of the transcripts differentially expressed in leaves of plants grown under SOIC with or without AA treatment (SOIC+AA vs. SOIC-AA, respectively) (see also **Supplemental Table 4**). Numbers of up- and down-regulated transcripts in each categorical group are indicated by pale and deep brown bars, respectively. Differentially expressed transcripts were sorted according to putative functional categories assigned by MapMan software. The figure was created using BioRender.com.

**Supplemental Figure 3: Climatic conditions of the experiment**. Maximum (max) and minimum (min) temperatures (A), relative humidity (B) and maximum photosynthetically active radiation (PAR; C) were recorded daily by two sensors.

**Supplemental Figure 4: Schematic representation of the experimental design.** At the beginning, all plants were irrigated under optimal irrigation conditions (OIC). When seedlings had developed 2-4 true leaves, they were transplanted into 17 L pots filled with a mixture of peat: coconut fibre: vermiculite (45:45:10), and grown under OIC. Seven days after transplanting, at the stage of 6-8 true leaves, tomato plants were divided in two groups: plants grown under OIC and plants grown under suboptimal irrigation conditions (SOIC, 60% of OIC irrigation). Applications of 20 mM AA were made weekly from the 2-4 true leaf developmental stage until the beginning of fruit collection. Leaf materials for biochemical and molecular characterizations were collected after 8th AA treatment and 30 days after water shortage, respectively.

**Supplemental Figure 5:** Volumetric content (A,B), soil temperature (T soil; C,D) and bulk electrical conductivity (BEC; E,F) under optimal irrigation conditions (OIC; A, C, E) and suboptimal irrigation conditions (SOIC; B, D and F) during the experiment.

