## Supplementary figures and images for "Acetic acid enhances tolerance to long-term water deficit in tomato by partially buffering transcriptomic and proteomic reprogramming independently of canonical jasmonate signalling"

### Supplemental Figure 1

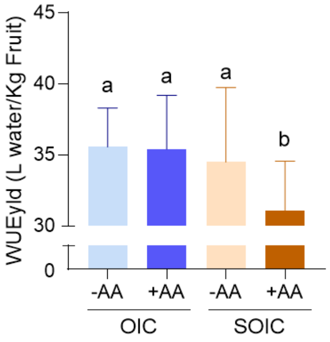


Supplemental Figure 1

### Supplemental Figure 2

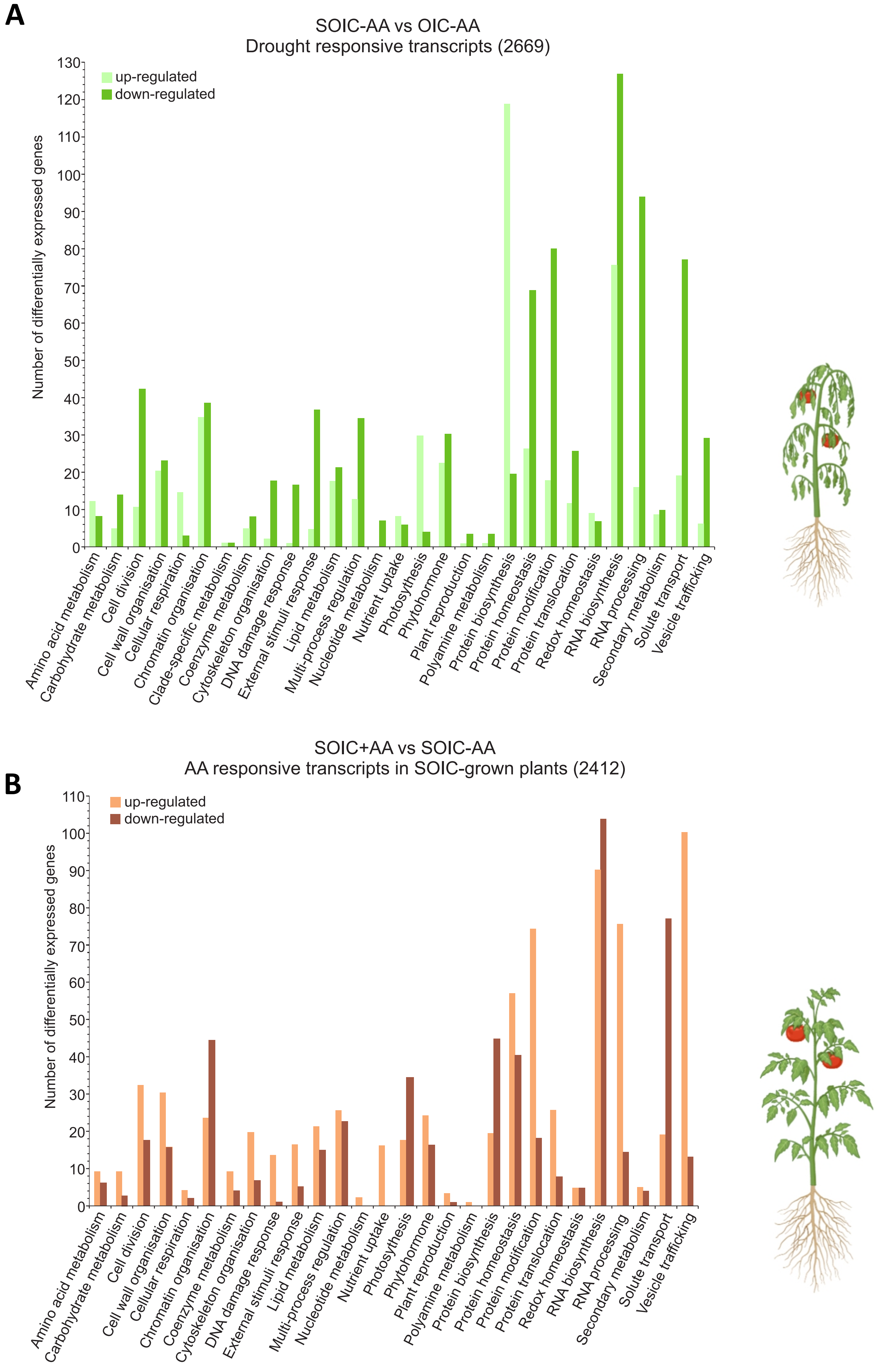


Supplemental Figure 2

### Supplemental Figure 3

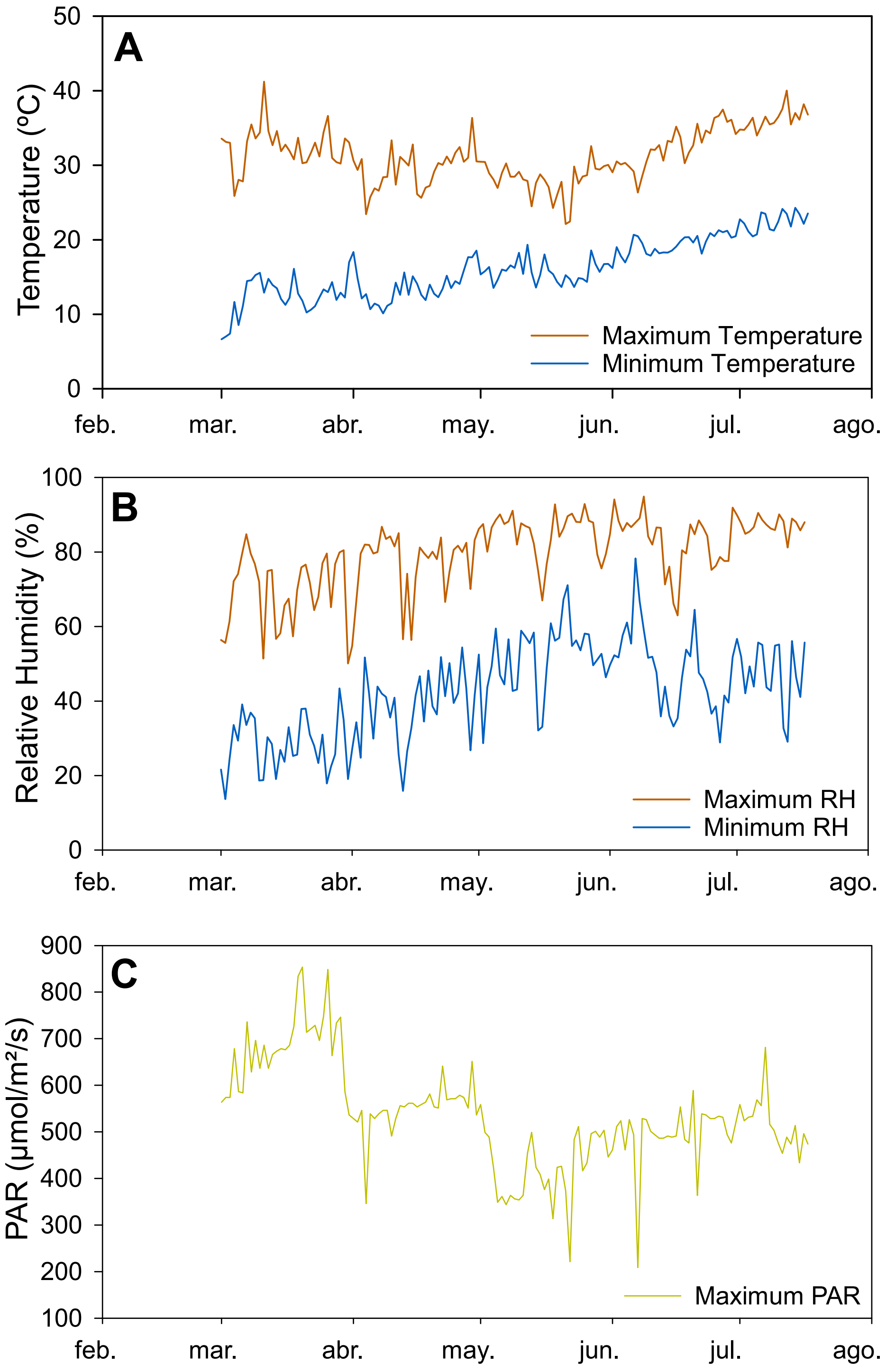


Supplemental Figure 3

### Supplemental Figure 4

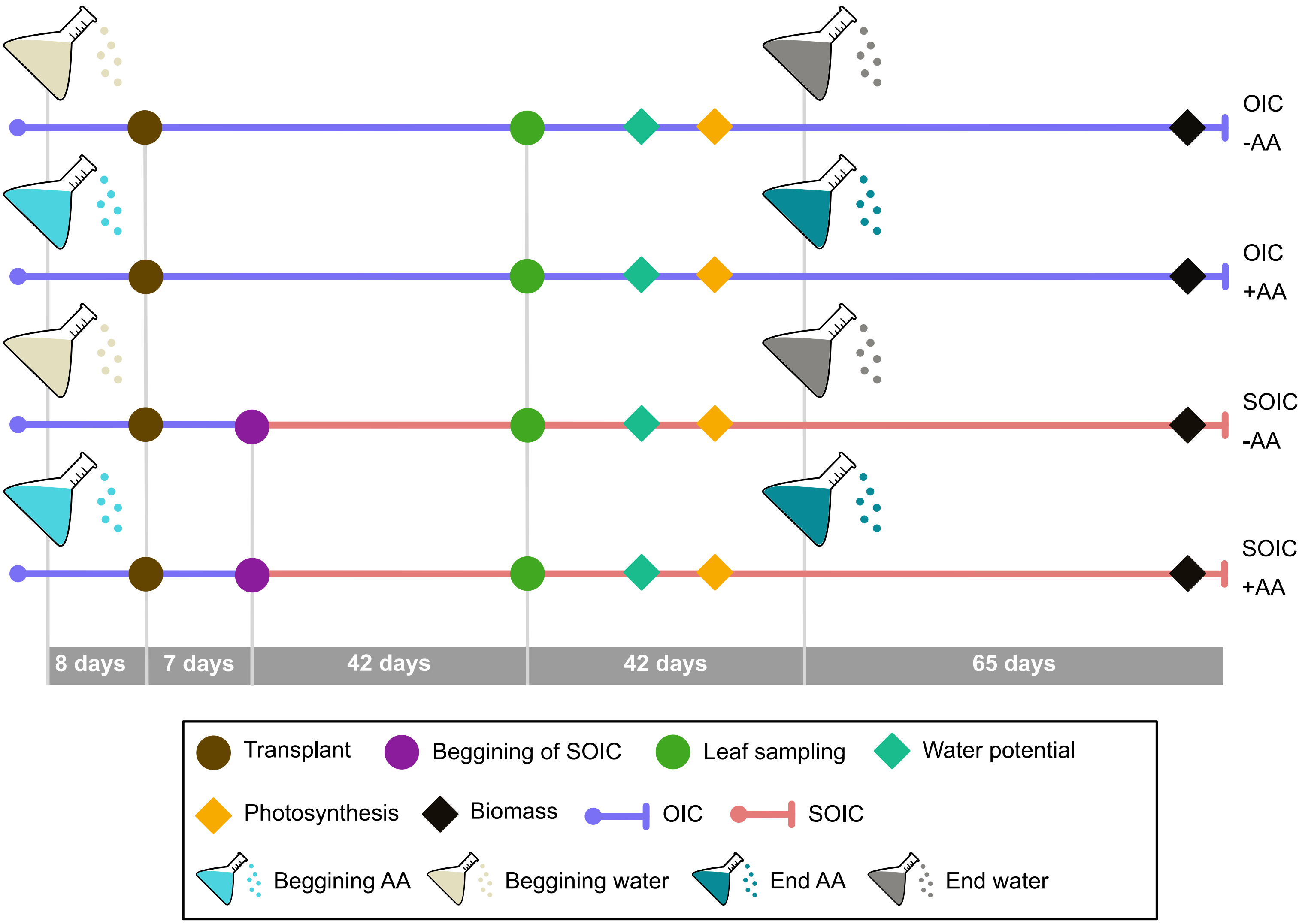


Supplemental Figure 4

### Supplemental Figure 5

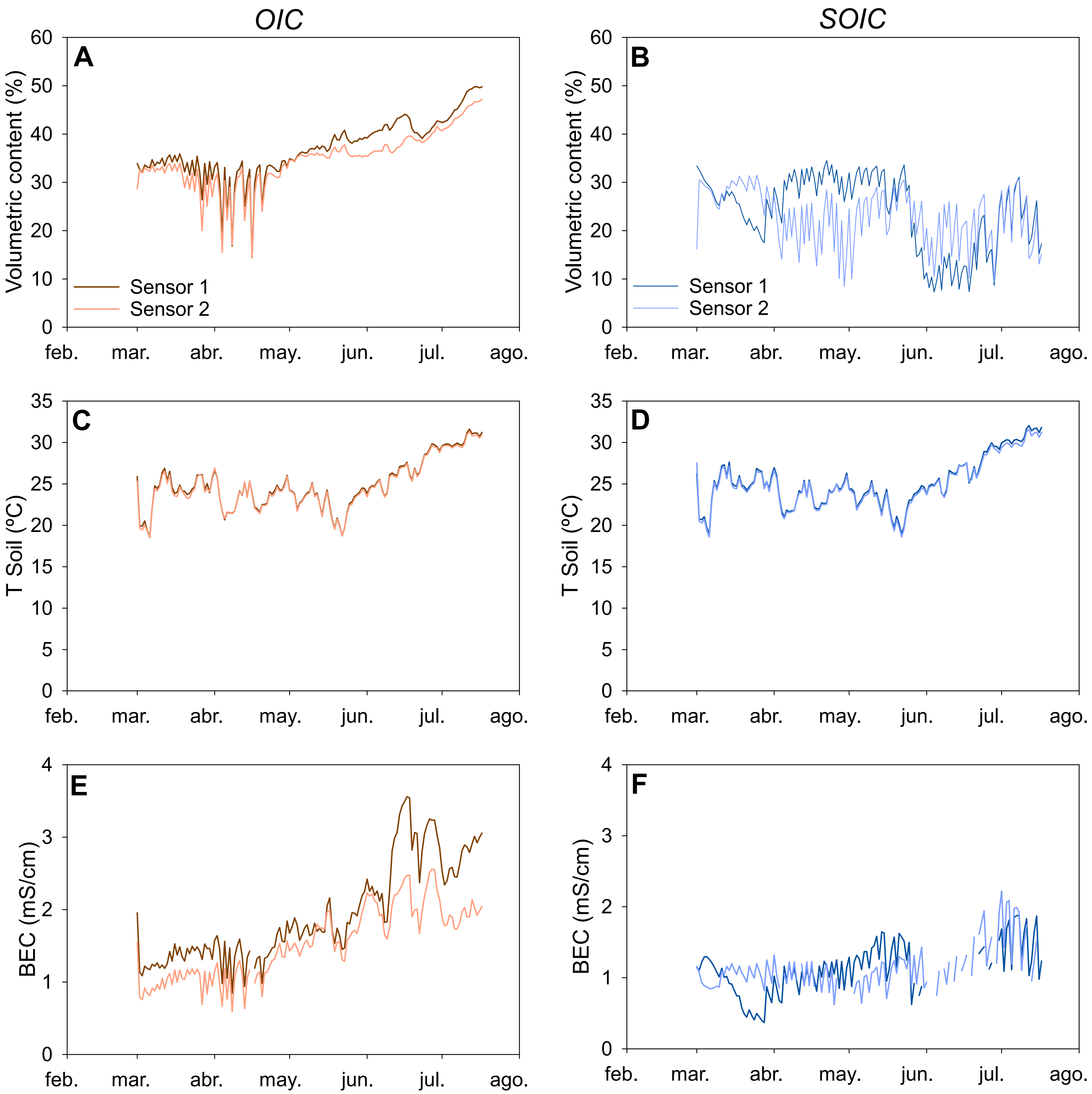


Supplemental Figure 5
